# Multi-sugar fermentation of lignocellulosic hydrolysate under industrially relevant conditions: comparison of several yeast strains

**DOI:** 10.1101/2022.08.12.503730

**Authors:** Allan Froehlich, Sean Covalla, Romain Fromanger, Joonas Hämäläinen, Tarja Kaartinen, Beth Mastel, M. Minna Laine

## Abstract

Both hexose- and pentose-fermenting yeasts are commercially available. The aim of this study was to test several yeast strains for their ability to ferment lignocellulosic feedstock and to evaluate their usability in a bioethanol production process based on the Cellunolix^®^ concept. The Cellunolix^®^ bioethanol demonstration plant of St1 uses sawdust to produce second-generation lignocellulosic ethanol. The study was performed in collaboration with yeast providers using two types of pretreated and filtered lignocellulosic ethanol hydrolysate originating from pine and willow. Ten pentose- and three hexose-fermenting yeast strains were tested. They all performed well under industrial conditions but differed in the rate of detoxification and profile for utilization of different sugars. A satisfactory 82–97% fermentation yield of a multi-sugar hydrolysate containing glucose, mannose, galactose, arabinose and xylose was achieved within 48 h. The results indicate significant potential for the usability of pentose-fermenting strains with real industrial hydrolysates and settings.

## 1. Introduction

St1 operates several small biorefineries in the Nordic countries that produce second-generation advanced fuel ethanol from surplus bread and bakery waste (Etanolix^®^), biowaste (Bionolix^®^) or sawdust (Cellunolix^®^). Pretreated softwood contains mainly sugars originating from cellulose, though some hemicellulose-derived sugars are also present in the Cellunolix^®^ feedstock.

Lignocellulosic substrates are composed of a wide variety of compounds—from different pentose and hexose sugars to organic acids and phenolic and furanic compounds [1–4]. The components present in industrial hydrolysates affect fermentation. A lag phase preceding fermentation is often caused by inhibitors, such as furanic compounds or acetic acid [2,5–8]. The presence of other substrates affects the sugar fermentation order and rate [9]. The laboratory experiments that compare xylose fermentation by different yeasts have often added glucose and xylose in a rich laboratory medium that contains all the necessary nutrient amendments [10] and, thus, have little correspondence to real-life situations in bioethanol production plants.

Robust and versatile yeast strains are required for satisfactory performance in industrial conditions. Metabolic engineering of bioethanol-producing yeasts strongly focuses on stress tolerance, xylose fermentation and acetic acid metabolism [11–14]. For example, acetic acid tolerance may be improved by altering yeast cell wall sphingolipids [15]. Inhibitor tolerance of the yeast can be further broadened by manipulating the redox-cofactor specificities of alcohol dehydrogenase [16,17].

As mentioned, acetic acid inhibits ethanol production [18]. If acetic acid could also be metabolized into ethanol, this would increase the production of ethanol. Strains that produce ethanol instead of glycerol in the presence or from acetic acid have been constructed by replacing glycerol-3-phosphate dehydrogenase (GPD1 and 2) with acetylating NAD-dependent acetaldehyde dehydrogenase [19] or acetylating aldehyde dehydrogenase (A-ALD) [16]. Thus, organic acids could be transformed into inducers or even co-substrates for ethanol production.

Carbon catabolite repression (CCR) affects the order in which yeast will utilize different sugars [20,21] and can be taken into account in the system—especially when the variety and amounts of different sugars cannot be pre-adjusted—by coupling gene expression under the same regulatory system or with genes promoting cell growth and biomass production, product accumulation, or both [22].

Polyploid yeasts have desirable properties for adaptation to stressful environments. It is possible to construct robust industrial strains by selecting a haploid strain with favored properties and crossing it with another haploid. However, controlled genetic alterations may be more difficult using polyploid strains [23–30].

In conclusion, in order to survive in an industrial-scale second-generation bioethanol plant, the fermenting yeast has to be able to adequately manage stress, be ferment multi-sugar hydrolysates, and cope with harsh conditions. Several components impact the fermentation performance as well as the fact that nutrients and other fermentation conditions are not necessarily optimal nor at constant levels at an industrial scale due to operability or cost-related matters.

Thus, usability evaluation should ideally test the strains under conditions that are as close as possible to those taking place in industrial surroundings.

In this study, the performance of recombinant pentose-fermenting yeasts was tested using real-life hydrolysates under industrially relevant conditions. More precisely, the fermentation experiments in the laboratory were performed under the same conditions that are in use in the current industrial-scale operation in the Cellunolix^®^ biorefinery.

## 2. Materials and methods

### 2.1. Strains

Eleven yeast strains capable of fermenting pentose and hexose sugars, Y52001–Y52011, were kindly provided by different yeast suppliers. Since some of the strains performed in a similar way during fermentation pre-testing, only the results from seven strains are shown, and data from three strains having only slight variations from several of the other strains are not presented. Three commercial and not genetically modified *Saccharomyces cerevisiae* strains capable of fermenting hexose sugars, Y63001–Y63003, were used as a reference. Strain Y52001 (used in the bioreactor experiment) is a strain further modified from Y52002 (used in the fermentation pre-test), where an antibiotic-resistance marker gene was removed.

### 2.2. Test hydrolysates

The fermentation performance of the yeast strains was tested using a hydrolysate originating from Cellunolix^®^, an industrial-scale bioethanol plant in Kajaani (Finland) that uses softwood sawdust as feedstock. The hydrolysate was pine sawdust that had been pretreated, enzymatically hydrolyzed and filter-pressed through a polypropylene-monofilament cloth filter (Hydrolysate 1, Table 1). In order to create a truly challenging environment for the fermenting yeast, the softwood hydrolysate was mixed with pretreated, filtered (in a funnel with suction using a Whatman paper) willow hydrolysate from laboratory-scale experiments to increase the amount of xylose and acetic acid in the medium (Hydrolysate 2, Table 1).

**Table 1.** Characteristics of the test hydrolysates. The softwood hydrolysate (Hydrolysate 1) was retrieved from an operating industrial-scale bioethanol plant. The willow hydrolysate was produced in laboratory-scale pretreatment and hydrolysis experiments. In Hydrolysate 2, these two aforementioned hydrolysates were mixed at a 75:25 (v/v) 1 ratio.

| Parameter (g/L) | Hydrolysate 1: Pretreated Pine Sawdust | Hydrolysate 2: Pretreated Pine Sawdust (75) and Willow (25) (v/v) Mixture |
| --- | --- | --- |
| Glucose | 53.04 | 59.16 |
| Arabinose | 1.82 | 1.30 |
| Galactose | 3.63 | 2.95 |
| Xylose | 6.32 | 10.38 |
| Mannose | 18.50 | 14.13 |
| Total fermentable sugars | 81.48 | 87.90 |
| Succinic acid | 0.58 | 1.02 |
| Acetic acid | 3.83 | 5.15 |
| Levulinic acid | 2.34 | 2.20 |
| Methanol | 0.14 | nd<sup>2</sup> |
| 5-hydroxymethylfurfural (HMF) | 2.62 | 1.77 |
| Furfural | 0.71 | 0.75 |
| NH<sub>4</sub>-N | 2.15 | na<sup>3</sup> |
| pH | 5.31 | 5.01 |

<sup>1</sup> v/v: volume/volume; <sup>2</sup> nd: not detected; <sup>3</sup> na: not analyzed.

### 2.3. Full-scale fermentation at the Cellunolix^®^ site

Industrial-scale fermentation at Kajaani was performed in a 350 m^3^ volume using the same pretreated softwood hydrolysate batch (Hydrolysate 1) that was used in the laboratory experiments. The yeast preparation containing hexose-fermenting yeast Y63001 was used with an inoculum size of 1.03 kg yeast cream per m^3^ fermentation volume, which corresponds to 0.33 g cell dry weight (CDW) per liter. Fermentation was performed in a fed-batch mode, and it was followed for 24 (batch phase started), 48, 72 and 98 h. Hydrolysate 1 was sampled from the filling line of the actual fermentation at site, so the same batch was fermented in the laboratory as in the full-scale plant.

### 2.4. Laboratory fermentation experiments

Tests were performed in shake flasks or serum bottles (fermentation pre-test) as well as in bioreactors under standardized conditions (Table 2). The pre-fermentation was followed for 48 or 72 h.

**Table 2.** Specifications of the laboratory tests. Results from the fermentation pre-tests in serum bottles and shake flasks were compared between each other, and results from the bioreactor experiments were compared with each other.

| Parameter | Serum bottle or shake flask tests = fermentation pre-tests |  |  | Bench-scale test in a bioreactor |  |
| --- | --- | --- | --- | --- | --- |
| Test vessel | Sealed 60 mL serum bottles | 250 mL shake flasks with fermentation water airlocks | 250 mL baffled shake flasks without airlocks | 3 L bioreactor | 2 L benchtop vessel |
| Test hydrolysate | Hydrolysate 1 | Hydrolysate 1 | Hydrolysate 1 | Hydrolysate 2 | Hydrolysate 2 |
| Test volume | 30 mL | 130 mL | 30 mL | 1830 mL | 1500 mL |
| Yeast propagation | YPD<sub>40</sub> medium containing 10 g/L yeast extract, 20 g/L peptone, 40 g/L glucose | YPD<sup>1</sup> medium containing 10 g/L yeast extract, 20 g/L peptone, 20 g/L glucose | Yeast extract–peptone–dextrose medium containing 20 g/L yeast extract, 10 g/L peptone, 100 g/L dextrose, and 1 mL/L 1:100 Lubrizol | YPD medium | Yeast extract–peptone–dextrose medium with Lubrizol |
| Yeast inoculum size in test volume | 0.5 g/L CDW<sup>2</sup> | $2 \times 10^7$ cells per mL ($A_{600}^3 = 0.7$) | $\sim 1$ g/L CDW ($OD_{600}^4 = 1.8$) | $2 \times 10^7$ cells per mL ($A_{600} = 0.7$) | $\sim 1$ g/L CDW ($OD_{600} = 1.8$) |
| Negative control | Media blank samples incubated and sampled under the same conditions as the inoculated bottles | Shake flask with no inoculum (i.e., sterile tap water instead of cell inoculum) | Abiotic flask incubated at 32 °C 150 rpm<sup>5</sup> for the duration of the experiment | Bioreactor with no inoculum (i.e., 1800 mL hydrolysate and 30 mL sterile tap water without cells) | Abiotic flask incubated at 32 °C 150 rpm for the duration of the experiment |
| Fermentation conditions | Duplicate bottles at 32 °C, 170 rpm, no added nutrients or pH control | Duplicate bottles at 32 °C, 120 rpm, no pH control, 5–6% (v/v)<sup>6</sup> YPD coming with inoculated cells | Duplicate bottles at 32 °C 150 rpm, no added nutrients or pH control | 32 °C, stirring at 30 rpm, no added nutrients or pH control | 32 °C, stirring at 200 rpm, no added nutrients or pH control, 1 mL/L 1:100 Lubrizol as antifoam agent |

<sup>1</sup> YPD: Yeast extract–Peptone–Dextrose medium; <sup>2</sup> CDW: Cell Dry Weight; <sup>3</sup> A<sub>600</sub>: Absorbance at 600 nm; <sup>4</sup> OD<sub>600</sub>: Optical Density at 600 nm; <sup>5</sup> rpm: Revolutions Per Minute; <sup>6</sup> v/v: volume/volume.

Additional testing using the same hydrolysate was performed in the premises of the yeast suppliers. Eleven strains were tested at the yeast provider’s laboratories with Hydrolysate 1 and two with Hydrolysate 2 in either shake flasks or serum bottles (fermentation pre-test). Additional chemical analyses in the end of the fermentation pre-test (t = 48 h) were performed from the samples sent to the St1 laboratory.

Two of the most prominent strains (or their modifications) were shipped to St1 Oy for further testing in bioreactors together with one pentose and one hexose-fermenting strain. In addition, one pentose and one hexose-fermenting strain were tested in a bioreactor-scale in the yeast supplier’s laboratory.

#### 2.4.1. Fermentation pre-tests: serum bottle fermentations in 60 ml

The yeast strains Y52002, Y52004, Y52005, Y52009, Y52011 and Y63002 were propagated in 55 mL of YPD40 medium (10 g/L yeast extract, 20 g/L peptone, and 40 g/L glucose filtered with a 0.22 µm bottle top filter). Cultures were grown in 250 mL baffled flasks with flask caps at 32 °C, 200 rpm (revolutions per minute), for approximately 24 h. CDW was determined using a Sartorius LMA (liquid mass analyzer) following pelleting and washing of cell pellets.

Fermentations were performed in sealed 60 mL serum bottles containing 30 mL of Hydrolysate 1. Duplicate bottles were inoculated at 0.5 g/L CDW, capped and crimped with gray butyl rubber stoppers. No nutrients or pH adjustments were made to the substrate (starting pH of ∼5.3). Bottles were incubated at 32 °C, 170 rpm and sampled for high-performance liquid chromatography (HPLC) analysis at 48 h. The media blank samples were also incubated and sampled under the same conditions as the inoculated bottles.

#### 2.4.2. Fermentation pre-tests: shake flask fermentations in 30 ml

Yeast strains Y52006 and Y63003 were first inoculated from a YPD (yeast extract–peptone– dextrose) plate to defined medium containing dextrose and grown for 8 h. Then, 30 mL of test Hydrolysate 1 was inoculated from the defined medium at OD600 (optical density at 600 nm) = 1, and grown at pH 5.3, 32 °C, 300 rpm, in 250 mL baffled shake flasks. The fermentation test was performed in duplicate on production shake flask with 1.0 g/L CDW pitch, 120 rpm at pH 5.3 with no airlock.

#### 2.4.3. Fermentation pre-tests: shake flask fermentations in 130 ml

Yeast strains Y52003 and Y63001 were pre-grown on 30 mL of YPD medium overnight (20–24 h), then transferred to 50 mL YPD over the weekend (70 h), and then transferred to 50 mL YPD overnight (20 h) at 32 °C, 150 rpm. The inoculation in the test volume was 2 × 10^7^ cells per mL. Yeast inocula were adjusted using the preconfigured cell growth method on the Thermo Scientific(tm) Evolution(tm) 60S UV–Visible Spectrophotometer, which estimates based on the relation where a value of A600 (absorbance at 600 nm) = 1 in growth medium corresponds to approximately 3 × 10^7^ yeast cells.

A 120 mL aliquot of Hydrolysate 1 was inoculated with the correct amount of yeast in growth medium, and the volume was made up to 130 mL with sterile tap water. No pH adjustment or addition of nutrients were carried out, although the inoculation in growth medium brought 5–6% (volume/volume) YPD into fermentation solution. The negative control was a shake flask with no inoculum (only sterile tap water added to the same reaction volume). The fermentation took place in duplicate 250 mL shake flasks with fermentation water airlocks at 32 °C, 120 rpm.

#### 2.4.4. Bioreactor experiments: bioreactor fermentations in 1.8 L volume

Bench-scale fermentation was performed in a three-liter bioreactor that was sterilized prior to adding hydrolysate, which was carried out in a biosafety cabinet. The hydrolysate that was a mixture of pine and willow hydrolysates (Hydrolysate 2, Table 1) was not sterilized. Inoculum was added to the ready-assembled reactors at the bench. The test volume was 1830 mL (1800 mL hydrolysate and 30 mL cell pellets in sterile tap water). Yeast strains Y52001, Y52003, Y52004 and Y63001 were propagated from freezer stocks in 30 mL YPD overnight (20–24 h), then transferred to 50 mL YPD over the weekend (67 h), and then transferred to 250 mL YPD overnight (20 h) at 32 °C, 150 rpm. Cells were harvested by centrifugation (4000 rpm, 2 min) and resuspended into sterile tap water before inoculation of the reactors. The inoculation in the fermentation volume was 2 × 10^7^ cells per mL (corresponding to A600 = 0.7). A bioreactor with no inoculum (i.e., 1800 mL hydrolysate and 30 mL sterile tap water without cells) was used as a negative control. Fermentation conditions were as follows: temperature 32 °C, stirring 30 rpm, and no added nutrients, aeration nor pH control. Sugars, ethanol, acetic acid, glycerol, furanic metabolites, pH, redox, oxygen, temperature were followed during the fermentation. The bioreactors were sampled aseptically at t = 0, 2, 4, 8, 10, 12, 21, 24, 44, 48, 52, 67, 72, and 141 h. The bioreactors had probes for temperature, pH and Redox measurements. The oxidation reduction potential (ORP) in bioreactors was measured using Mettler Toledo Pt4805-SC-DPAS-K8S/225 combination Redox electrode. The pH was measured using EasyFerm Plus Arc 225 electrode from Hamilton Inc., and dissolved oxygen using VisiFerm DO Arc 225 electrode, also from Hamilton Inc.

#### 2.4.5. Bioreactor experiments: bioreactor fermentations in 1.5 L volume

Additional testing using the same hydrolysates was performed in the premises of the yeast supplier. Yeast strains Y52006 and Y63003 were propagated in 30 mL of a medium containing 20 g/L yeast extract, 10 g/L peptone, 100 g/L dextrose, and 1 mL/L 1:100 diluted Lubrizol. Cultures were grown in 250 mL baffled flasks at 32 °C, pH 5.5, 300 rpm for 6 h. The inoculum was then transferred to 100 mL test hydrolysate with nutrients and Lubrizol for overnight. The cells were harvested by centrifuge (4000 rpm, 1 min), supernatant was decanted, and the pellet resuspended in assay medium at ∼1 g/L CDW (OD600 = 1.8). As a negative control, an abiotic flask was incubated at 32 °C, 150 rpm, for the duration of the experiment.

Two-liter bench fermentations were conducted in blended willow and pine Hydrolysate 2. The seed for the production vessels was cultivated in a two-step shake flask protocol started from a frozen stock of yeast provider’s culture collection. The stock was plated onto a YPD plate and allowed to grow overnight at room temperature, then used to inoculate 30 mL YPD liquid media that was placed in a 250 mL baffled shake flask at pH 5.5, 32 °C, 300 rpm, for 6 h. The seed culture for the batch fermentation was inoculated to 0.3 g/L in 100 mL of 100% Hydrolysate 2 in a 500 mL baffled shake flask supplemented with 20 g/L yeast extract and 10 g/L peptone. The pH of the flask was not altered and was used as received at pH 5.3. The culture was allowed to grow at 32 °C for 16 h at an agitation of 300 rpm. To control any possible foaming during the fermentation, a dilute antifoam solution was added (1 mL/L of 1:100 Lubrizol BCC 627 defoamer).

The 2 L stirred tank batch fermentations were inoculated from seed harvested from active shake flasks targeting strains Y52006 and Y63003 at 1.0 g/L CDW, or OD600 = 1.8. The 2 L production vessels were run at 1.5 L working volume at temperature of 32 °C using Hydrolysate 2, with no additional nutrients added. The pH of the hydrolysate was not adjusted or controlled in any way during the fermentation run. The stirred tanks were agitated at 200 rpm. A dilute antifoam solution was added (1 mL/L of 1:100 Lubrizol), to prevent any possible foaming during the fermentation.

### 2.5. Chemical measurements

#### 2.5.1. Analysis of furanic compounds

Furanic metabolites were analyzed by the HPLC method further developed from the one published by Yuan [31] using the Rezex ROA 300 mm × 7.8 mm column. The eluent was acetonitrile/5 mM sulfuric acid (17:83 vol%) at flow rate of 0.5 mL/min. Column oven temperature was 25 °C. The compounds were detected using UV (ultraviolet) detector at 30 °C, at both 215 and 254 nm. Hydroxymethylfurfural (HMF, 5-(hydroxymethyl)furan-2-carbaldehyde) alcohol was run as a separate standard diluted into methanol. The detection limits for different furanic compounds were as follows (measurement wavelength in parentheses): HMF acid (254 nm), 0.006–0.50 g/L; HMF alcohol (215 nm), 0.044-2.21 g/L; HMF (5-(hydroxymethyl)furan-2-carbaldehyde, 254 nm), 0.016– 3.13 g/L; 2-furoic acid (215 nm), 0.006–1.26 g/L; furfuryl alcohol (215 nm), 0.012–1.60 g/L, and furfural (215 nm), 0.013–2.57 g/L.

#### 2.5.2. Analysis of sugars, organic acids and ethanol

Sugars were analyzed by ion chromatography using Dionex CarboPac SA10 (4 mm) column with guard column CarboPac SA10G. The eluent was 1 mM KOH at 1.5 mL/min, 45 °C. The sugars were detected using Integrated Amperometry, Carbohydrate Standard Quad Waveform with PTFE Gold as (disposable) working electrode and silver/silver chloride as a reference electrode. The detection range for different sugar components was, depending on the compound, from 12 to 250 µg/mL. The sugar compounds detected were D-cellobiose, D(+)-glucose, L(+)-arabinose, D(+)-galactose, D(+)-mannose, D(+)-xylose, fructose and sucrose.

Organic acids, glucose and ethanol were analyzed using the same HPLC–UV/RI equipment with RI (refractive index) detection and Rezex ROA column as with the analysis of furanic compounds, except that the eluent was 5 mM H2SO4 with a flow rate of 0.6 mL/min and oven temperature of 60 °C. In the full-scale fermentation event, the chemical measurements were performed in the Cellunolix^®^ bioethanol plant’s laboratory where glucose was separated in an HPLC run, but the rest of the sugars were co-eluting with mannose. The equipment and column were similar to the abovementioned.

For the retained samples from 30 mL shake flask and 1.5 L reactor experiments, a standard Aminex 87H column was used with dilute H2SO4 as mobile phase, and RI and UV for detection [32]. For shake flasks without airlocks, the HPLC ethanol values were corrected for evaporation using an experimentally determined correction factor of 0.003.

### 2.6. Ethanol yield calculation

If not otherwise stated, the ethanol yield was calculated based on the sum of five fermentable sugars (glucose, galactose, arabinose, mannose and xylose) in the beginning of the fermentation. A factor of 0.511 (g ethanol/g sugar) was used to calculate the maximum theoretical conversion of sugars into ethanol.

## 3. Results and discussion

### 3.1. Industrial-scale fermentation

The hexose-fermenting strain currently being used in the industrial-scale Cellunolix^®^ plant ferments glucose and mannose into ethanol in 48 h (Figure 1). The ethanol yield was approximately 87% of the theoretical maximum of the two sugars. The ethanol formation started after 24 h when most of the furfural had been detoxified (Figure 1).

**Figure 1.**
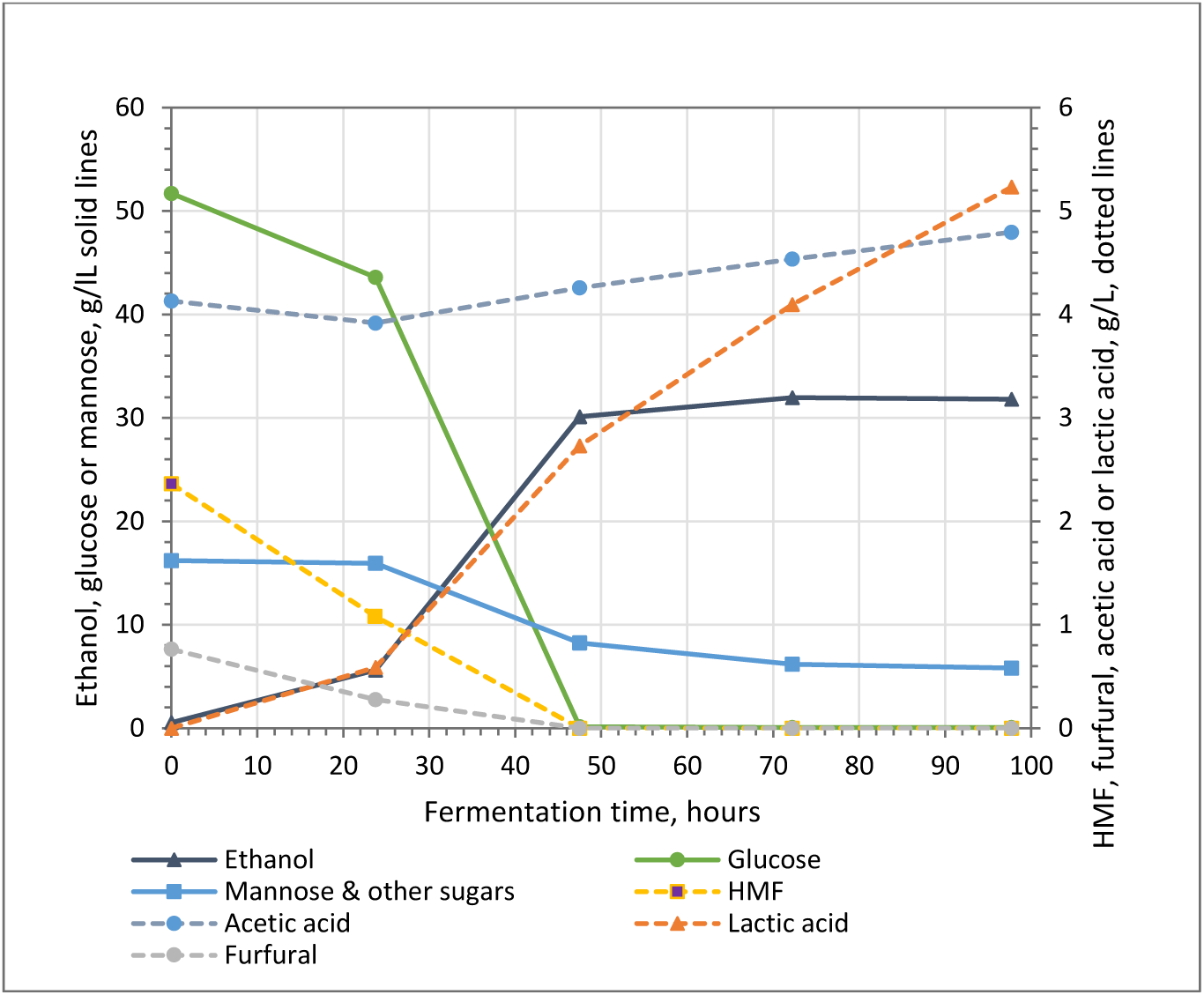
Industrial-scale fermentation using the same pretreated softwood hydrolysate batch (Hydrolysate 1) that was used in the laboratory experiments comparing yeast strains. The hexose-fermenting Y63001 strain was used in the fermentation with an inoculum size of 1.03 kg/m^3^ wet weight. The chemical measurements were performed in the laboratory of the Cellunolix^®^ bioethanol plant, where glucose is separated in a HPLC run, but the rest of the sugars are co-eluting with mannose.

### 3.2. Small-scale preliminary testing of fermentation at 48 h

The substrate utilization pattern of ten pentose- and three hexose-fermenting yeast strains was first screened in a small-scale laboratory test using shake flasks or serum bottles (Table 2). The same Hydrolysate 1 originating from the industrial-scale plant was used in the laboratory-scale fermentation experiments to evaluate the pentose-fermenting yeast strains (Table 1). The evaluation time, 48 h, was consistent with the fermentation time of the same hydrolysate in industrial scale (Figure 1).

The strain Y52004 had the broadest spectrum of fermentable sugars: it was able to ferment all five measured sugars present in the softwood hydrolysate: glucose, mannose, galactose, xylose and arabinose (Figure 2A, Table 3). It was the only strain among those tested that was able to utilize arabinose. Galactose was utilized by the strains Y52002, Y52004, Y63001 and Y63003. All pentose-fermenting strains fermented xylose, although it took a longer time of 94 h for the strain Y52006 to use 82% of available xylose under the test conditions (data not shown).

**Table 3.** Substrate utilization by the yeast strains during a 48 h fermentation pre-test. Closed circles = more than 90% of the scheme at 48 h, open circles = more than 50% of the substrate was utilized within 48 h; dash = less than 50% or none of the substrate was utilized.

| Strain | Acetic acid | Arabinose | Galactose | Glucose | Xylose | Mannose | Ethanol yield % | Ethanol, g/L |
| --- | --- | --- | --- | --- | --- | --- | --- | --- |
| Y52002 | o | - | ● | ● | ● | ● | 97 | 38.5 |
| Y52003 | - | - | - | ● | ● | ● | 86 | 32.8 |
| Y52004 | - | ● | ● | ● | ● | ● | 98 | 39.1 |
| Y52005 | o | - | - | ● | ● | ● | 94 | 37.4 |
| Y52006 | - | - | - | ● | o | ● | 92 | 37.8 |
| Y52009 | - | - | - | ● | ● | ● | 89 | 35.4 |
| Y52011 | - | - | - | ● | ● | ● | 90 | 35.8 |
| Y63001 | - | - | ● | ● | - | ● | 84 | 32.0 |
| Y63002 | - | - | - | ● | - | ● | 82 | 32.7 |
| Y63003 | - | - | ●/o | ● | - | ● | 88 | 32.5 |

**Figure 2.**
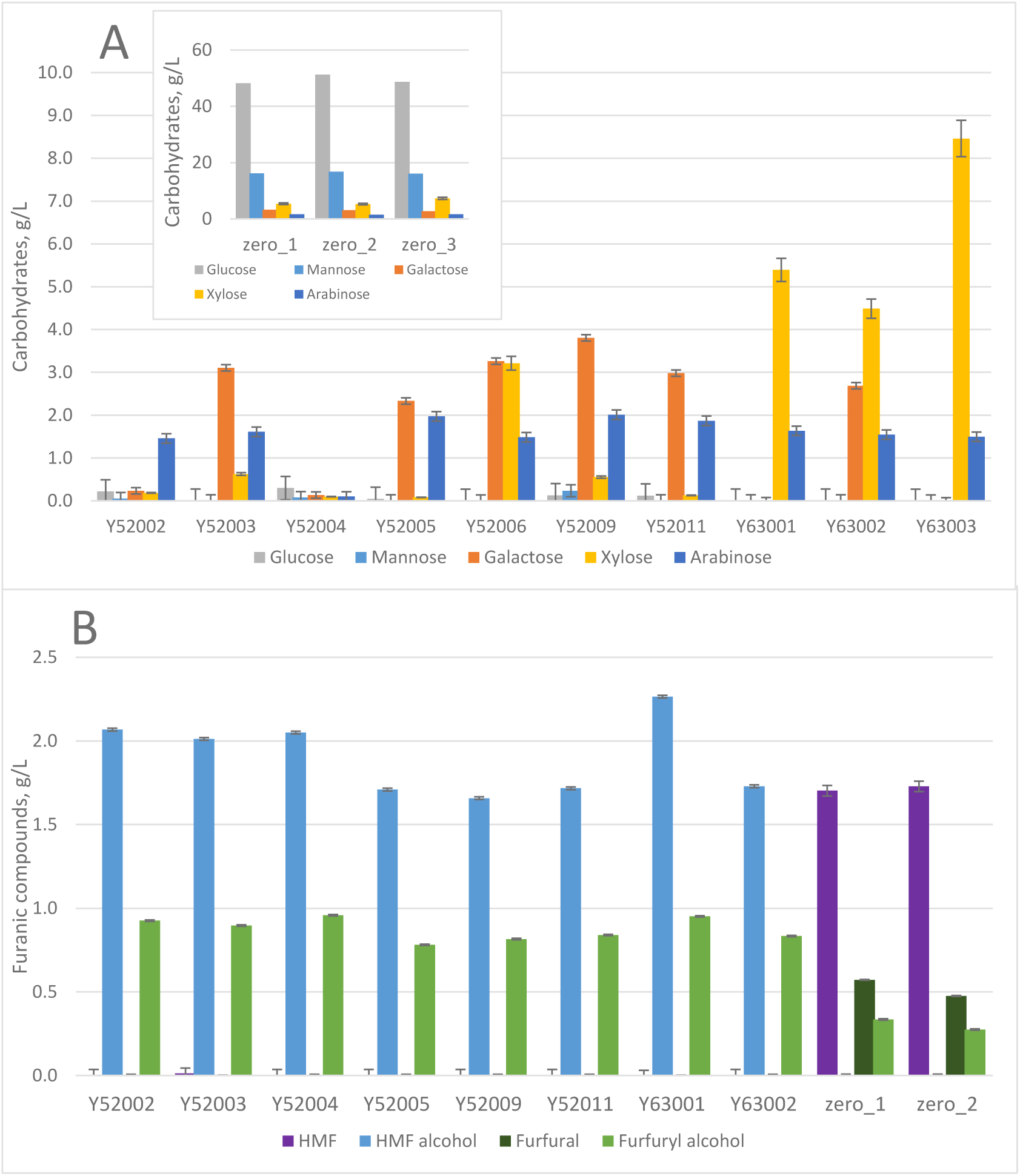
**(A)** Concentrations of different sugars at the 48 h timepoint in pre-fermentation experiments with Hydrolysate 1 using different yeast strains. **(B)** Products of transformation of furanic inhibitors into corresponding alcohols by the test strains at 48 h. Furanic compounds were not measured from the fermentation of strain Y52006.

All tested strains were able to transform furfural and HMF into furfuryl alcohol and HMF alcohol within 48 h when fermenting the softwood Hydrolysate 1 (Figure 2B).

### 3.3. Fermentation experiments with the four most prominent strains

#### 3.3.1. Fermentation performance in mixed test hydrolysate

Strains Y52001 (slightly modified from strain Y52002), Y52003, Y52004 and Y52006 were further studied in bioreactors with Hydrolysate 2. These strains were selected based on their broad spectrum of potential substrates and their ethanol yield produced from the softwood Hydrolysate 1 (Table 3, Figure 2A). Strain Y63001 was included in the experiments as a reference hexose-fermenting strain.

The softwood and willow hydrolysates were mixed to produce a Hydrolysate 2 (Table 1), where inhibitors such as furanic compounds and acetic acid appeared at concentrations slightly below the threshold values for *Saccharomyces cerevisiae* strains. The threshold concentrations were established in previous unpublished investigations carried out by St1.

The ethanol fermentation specificities and rates of the four selected strains were compared to Y63001 that was used as a reference hexose-fermenting yeast. The fermentation was followed up to 67 h. All pentose-fermenting strains performed well, with close to 90% ethanol yield achieved in 67 h. Strain Y52001 had the highest ethanol titer of 38 g/L, already after 48 h of fermentation. It was also able to utilize acetic acid unlike other tested strains (Table 4). Strain Y52004 had the slowest fermentation rate, most likely due to the slower detoxification rate of furfural (24–44 h) and 5-hydroxymethylfurfural (HMF, 44–48 h), but it reached the same final ethanol titer of 38 g/L within 67 h as the strain Y52001. Due to small differences in bioreactor setups, the initial concentration of fermentable sugars was smaller for the strain Y52006, so the ethanol titer was also smaller. Nevertheless, the ethanol recovery of the strain Y52006 was equally good in 67 h as with the rest of the tested strains (Table 4). The performance of the reference strain, hexose-fermenting Y63001, was exceptional, even though it could not ferment pentose sugars nor utilize acetic acid (Table 4).

**Table 4.**
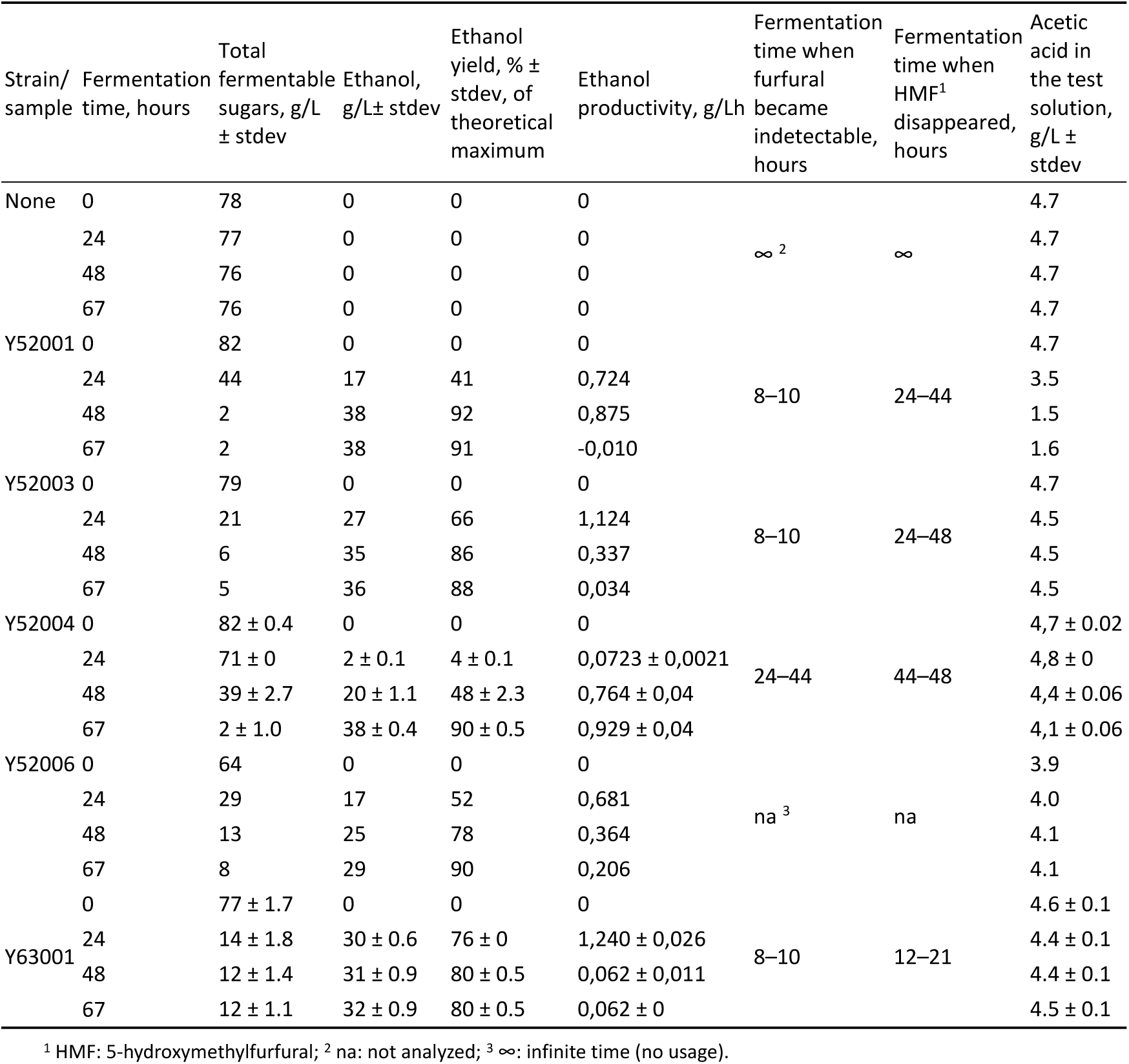
Comparison of fermentation performance by four pentose-fermenting strains tested in bioreactors with Hydrolysate 2. The results were compared to one hexose-fermenting yeast strain. The exact time point when furanic compounds were fully converted lies in between the sampling times shown here. Total fermentable sugars included glucose, mannose, galactose, arabinose and xylose. Hydrolysate 2 used was a mixture of filtered pine sawdust and willow hydrolysates. The differences in the total fermentable sugar concentration in the beginning of the test reactors are due to small variations in the reactor setups. Strains Y52004 and Y63001 were run on duplicate reactors.

The fastest detoxifiers among pentose-fermenting strains in 8–10 h were Y52001 and Y52003, but the fastest of all tested strains was the hexose-fermenting reference strain Y63001. The reference strain was able to transform all furfural into furfuryl alcohol in less than eight hours and HMF into HMF alcohol in less than 21 h (Table 4, Figure 3).

**Figure 3.**
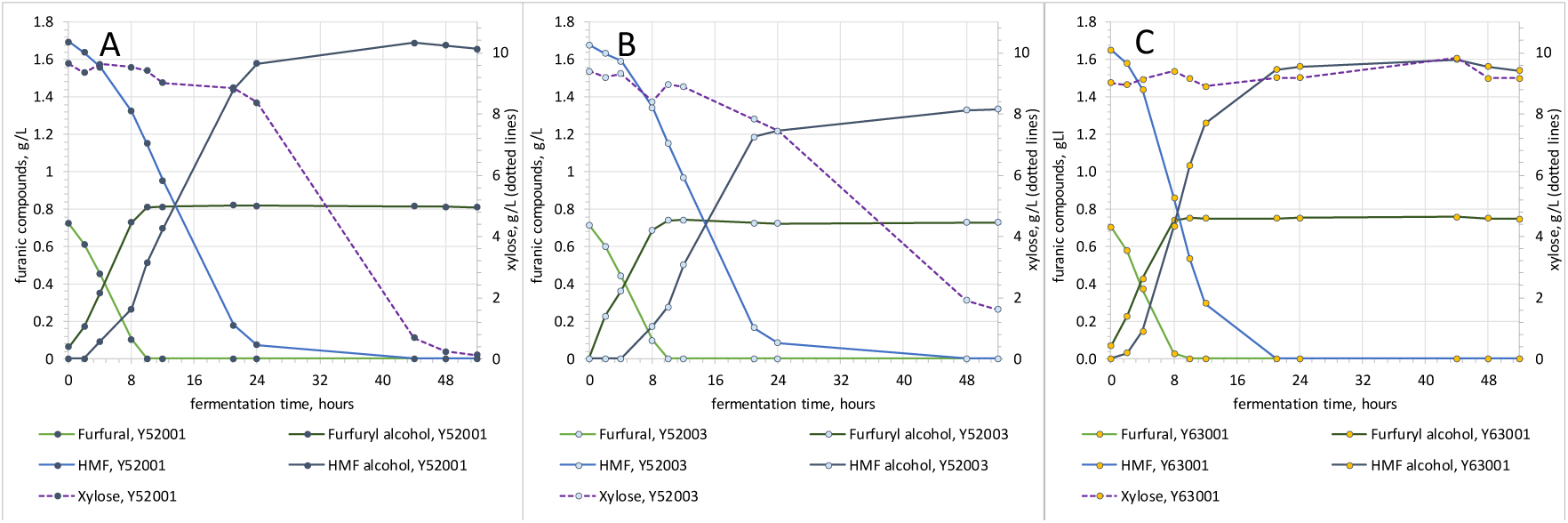
A closer look at inhibitor detoxification during bioreactor fermentation experiments in two pentose- and one hexose-fermenting yeast strains, Y52001 (A), Y52003 (B) and Y63001 (C). Detoxification of furfural and HMF into corresponding alcohols (left axis) are compared to the rate of xylose usage (axis on the right, dotted lines).

#### 3.3.2. Detoxification behavior

After the initial experiments, the two fastest-fermenting strains, Y52001 and Y52003, were further investigated in bioreactors with respect to the hexose-fermenting strain Y63001, and their detoxification behavior under the given circumstances was compared (Figure 3).

Furfural detoxification occurred at the same pace and extent among the three strains, but there was a difference in HMF metabolism between Y63001 and the two pentose-fermenting strains Y52001 and Y52003 (Figure 3). The hexose-fermenting strain Y63001 turned HMF into the corresponding alcohol faster than the pentose-fermenting yeast strains Y52001 and Y52003. The capability to ferment xylose seemed to result in slower HMF detoxification for these two strains. In addition, these two events followed each other, i.e., after HMF had been detoxified, xylose fermentation began (Figure 3). In fact, it has been reported that lignocellulose inhibitors affect more xylose than glucose fermentation [12,33,34]. HMF detoxification and xylose fermentation seem to be linked via NADPH or NADH usage [10,14,35,36].

The amount of acetic acid was relatively high, more than 4 g/L (68 mM) in the tested hydrolysate solutions (Tables 1 and 4). Acetic acid can affect the detoxification and fermentation in several ways. Small amounts of acetic acid (20 mM) can improve the HMF tolerance of the yeast [37]. Higher amounts (125 and 250 mM) of acetic acid clearly hinder the fermentation of glucose and xylose. There are differences in acetic acid tolerance between different yeast strains [25]. The undissociated form of acetic acid is inhibitory to yeasts; therefore, increasing the medium pH can mitigate the negative effect [38].

#### 3.3.3. Preferred order of sugar utilization

The specific usage of different sugars by the selected yeast strains was further investigated. Four out of five studied strains utilized glucose first, followed by xylose (if used) and/or mannose and, finally, the less frequently utilized arabinose and galactose, if they were utilizable by the strain (Table 5). The order of glucose > mannose > galactose utilizability follows the degree of respiration that *Saccharomyces cerevisiae* displays to different carbon sources [39].

**Table 5.**
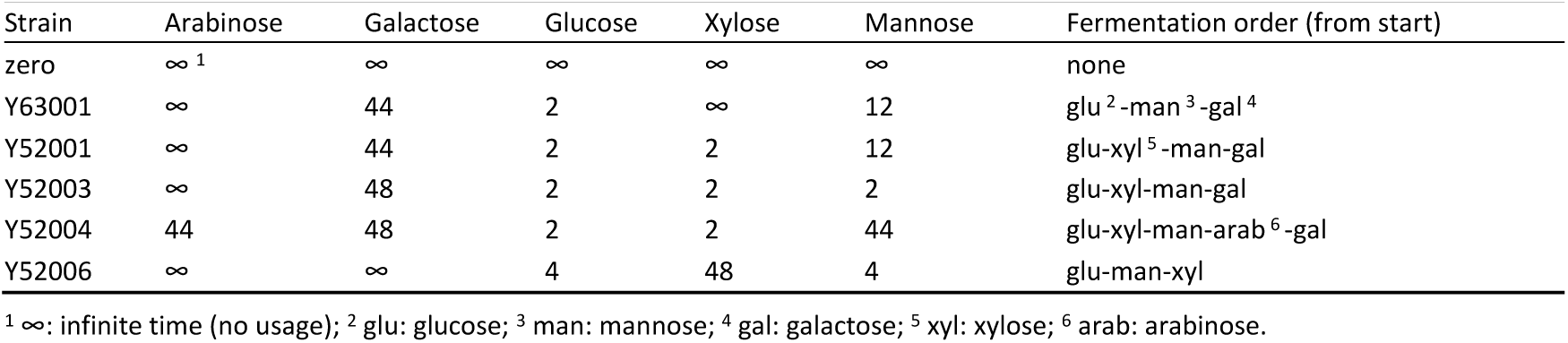
Preferred fermentation order of different sugars by the strains Y63001, Y52001, Y52003, Y52004 and Y52006. Numbers refer to the time point when the consumption of each sugar started.

| Strain | Arabinose | Galactose | Glucose | Xylose | Mannose | Fermentation order (from start) |
| --- | --- | --- | --- | --- | --- | --- |
| zero | ∞<sup>1</sup> | ∞ | ∞ | ∞ | ∞ | none |
| Y63001 | ∞ | 44 | 2 | ∞ | 12 | glu<sup>2</sup>-man<sup>3</sup>-gal<sup>4</sup> |
| Y52001 | ∞ | 44 | 2 | 2 | 12 | glu-xyl<sup>5</sup>-man-gal |
| Y52003 | ∞ | 48 | 2 | 2 | 2 | glu-xyl-man-gal |
| Y52004 | 44 | 48 | 2 | 2 | 44 | glu-xyl-man-arab<sup>6</sup>-gal |
| Y52006 | ∞ | ∞ | 4 | 48 | 4 | glu-man-xyl |

<sup>1</sup> ∞ : infinite time (no usage); <sup>2</sup> glu: glucose; <sup>3</sup> man: mannose; <sup>4</sup> gal: galactose; <sup>5</sup> xyl: xylose; <sup>6</sup> arab: arabinose.

Strain Y52006 had slightly different order of preference (Table 5) than rest of the compared strains since it fermented both glucose and mannose before xylose (Table 5).

#### 3.3.4. Redox and pH changes during fermentation

In the beginning, redox values decreased sharply, apparently due to the yeast using up the oxygen during the aerobic growth phase (Figure 4). Redox value continued to decrease while sugars, especially glucose, were reduced. Once the hexose fermentation was finished, the redox values began to increase again (Figure 4).

**Figure 4.**
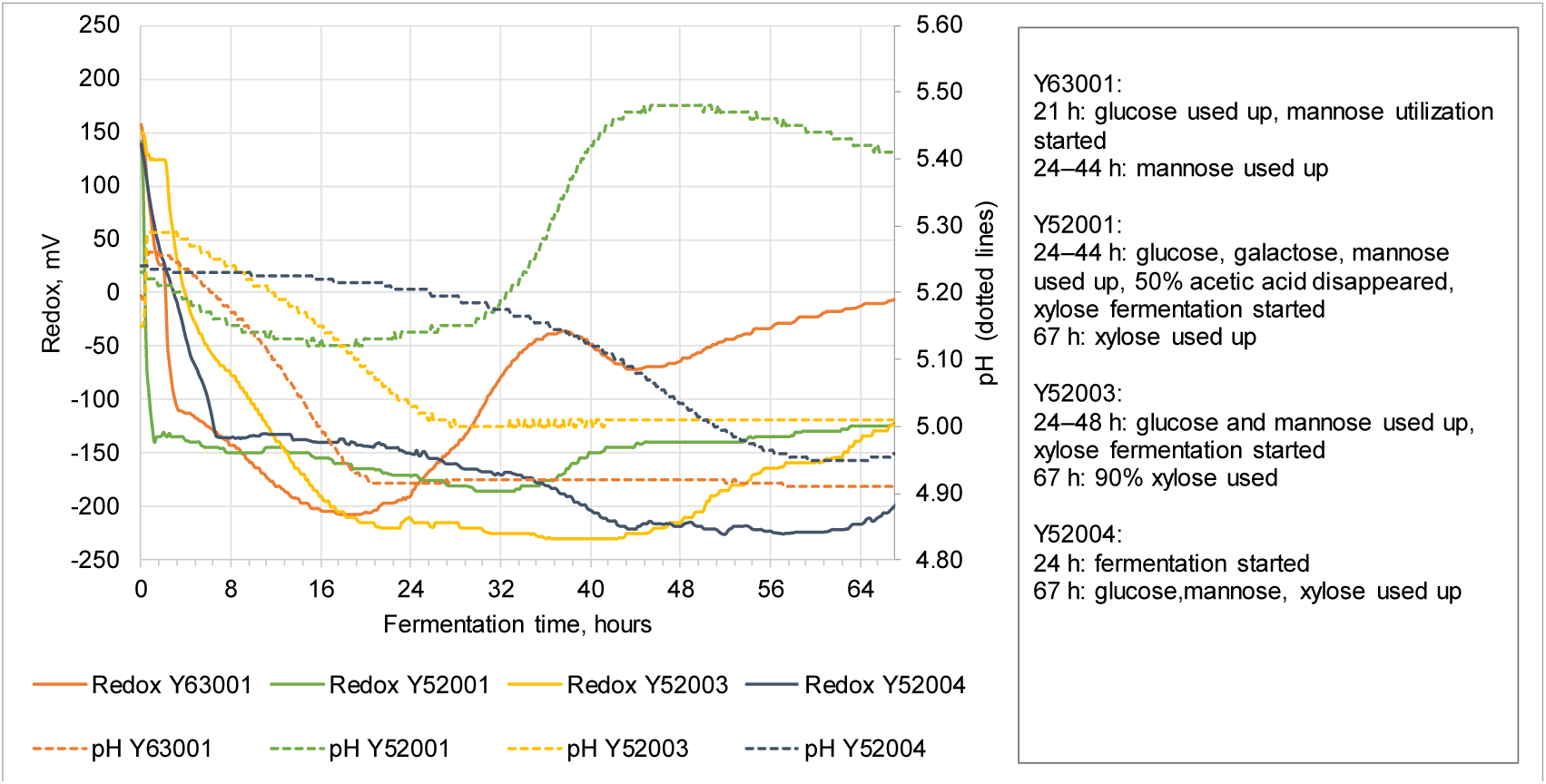
Redox and pH values of the three pentose- and one hexose-fermenting yeast strains, Y52001, Y52003, Y52004 and Y63001, during the bioreactor fermentation test. Time ranges of different events listed on the right reflect the sampling point.

The pH values decreased from around 5.2 close to 4.9 during the fermentation. The strain Y52001 was able to utilize acetic acid. Acetic acid was utilized in 24–44 h, and from 33 h onward, the pH increased to 5.4–5.5 (Figure 4).

Thus, redox and pH changed during the fermentation as the sugars were utilized, inhibitors detoxified and ethanol produced. During ethanol fermentation, the redox value changes due to cell growth when NAD(P)H/NAD(P)+ is used either in glycolysis or for biomass growth [40]. The redox value also reflects the amount of dissolved oxygen in the bioreactor, i.e., the presence of electron donors in the solution [40].

## 4. Conclusions

All studied strains performed well in terms of fermentation, even without any optimization (nutrients, oxygen, pH adjustment) of conditions. The choice of the most utilizable yeast strain depends on the nature of the hydrolysate: the ratio of different pentose and hexose sugars as well as the quality and quantity of the inhibitors present, and the required detoxification rate and time for the yeast under the conditions of the intended industrial-scale process. For industrial application, conditions could be optimized in order to improve the performance of all strains.

## Acknowledgments

We thank Hannu Kalliomäki, Minna Yamamoto and Jukka Lommi for performing pretreatment pilot tests for willow hydrolysate and Paula Aerikkala for laboratory assistance. We also thank Anna Kankaanpää for proof-reading the manuscript.

## Author Contributions

Conceptualization, M.M.L.; methodology, T.K.; validation, T.K.; investigation, A.F., S.C., J.H., B.M. and M.M.L.; resources, A.F., S.C., R.F., J.H., T.K. and B.M.; writing—original draft preparation, A.F, S.C., J.H., B.M. and M.M.L.; writing—review and editing, A.F., S.C., R.F, B.M., J.H. and M.M.L. All authors have read and agreed to the published version of the manuscript

